# Core-shell DNA-cholesterol nanoparticles exert lysosomolytic activity in African trypanosomes

**DOI:** 10.1101/2022.07.18.500428

**Authors:** Robert Knieß, Wolf-Matthias Leeder, Paul Reißig, Felix Klaus Geyer, H. Ulrich Göringer

**Affiliations:** Molecular Genetics, Technical University Darmstadt, 64287 Darmstadt, Germany

**Keywords:** DNA nanoparticles, African trypanosomiasis, drug design, lysosomal collapse, DNAzyme, dual-target, chemotherapeutic

## Abstract

*Trypanosoma brucei* is the causal infectious agent of African trypanosomiasis in humans and Nagana in livestock. Both diseases are currently treated with a small number of chemotherapeutics, which are hampered by a variety of limitations reaching from efficacy and toxicity complications to drug-resistance problems. Here, we explore the forward design of a new class of synthetic trypanocides based on nanostructured, core-shell DNA-lipid particles. In aqueous solution, the particles self-assemble into micelle-type structures consisting of a solvent-exposed, hydrophilic DNA shell and a hydrophobic lipid core. DNA-lipid nanoparticles have membrane-adhesive qualities and can permeabilize lipid membranes. We report the synthesis of DNA-cholesterol nanoparticles, which specifically subvert the membrane integrity of the *T. brucei* lysosome, killing the parasite with nanomolar potencies. Furthermore, we provide an example of the programmability of the nanoparticles. By functionalizing the DNA shell with a spliced leader (SL)-RNA-specific DNAzyme, we target a second trypanosome-specific pathway (dual-target approach). The DNAzyme provides a backup to counteract the recovery of compromised parasites, which reduces the risk of developing drug resistance.

## Introduction

The inherent complexity of developing, testing, and synthesizing new anti-infectives and therapeutics has never been more obvious than during the SARS-CoV2 pandemic. While modern-era drug development has benefitted immensely from the large amount of genome, proteome, and high-throughput data, the need for new and improved antiviral, antibacterial, antifungal, and antiparasitic medications has not vanished, and the search for alternative drug concepts and new drug targets has remained a key challenge in academic and industrial pharmaceutical research alike. This is especially the case for illnesses with unmet medical and therapeutic needs such as neglected tropical diseases (NTDs) (Feasey et al., 2010; Molyneux et al., 2018) including onchocerciasis, schistosomiasis, lymphatic filariasis, and human African trypanosomiasis (HAT). HAT, also known as African sleeping sickness, is caused by the two protozoan parasite species, *Trypanosoma brucei rhodesiense* and *Trypanosoma brucei gambiense*. While the incidence of HAT is currently at an unprecedented low (Büscher et al., 2017), large parts of Africa are still threatened by the disease (Kennedy, 2019). This is further aggravated by the zoonotic trypanosome species *T. congolense, T. vivax*, and *T. brucei brucei*, which thwart almost every agricultural progress in sub-Saharan Africa. HAT is fatal if left untreated, and due to the variable glycoprotein surface (VSG) of the parasite, all vaccination efforts have failed. As a consequence, the disease is treated by chemotherapy (Dickie et al., 2020). Unfortunately, only a small number of clinically relevant compounds are available and these compounds suffer in part from grave side effects, narrow therapeutic windows, and the problem of parenteral administration. The situation is further vexed by the occurrence of treatment failures due to the rise of drug-resistant parasite strains (Fairlamb and Horn, 2018). Consequently, despite the recent addition of fexinidazole as the first oral monotherapy against *T. b. gambiense* infections (Pollastri, 2018; Dickie et al., 2020), new and improved drugs are still needed for the treatment of HAT.

Here, we present an alternative drug development concept, which we term dual-target approach (Figure 1). The method specifically aims to reduce the incidence of drug resistance by targeting two parasite-specific biochemical pathways simultaneously. As the first target, we focus on the lysosomal membrane of *T. brucei*. Lipid membranes represent a core characteristic in all living systems, and multiple synthetic and biological compounds capable of disrupting lipid membranes have been identified (Regen, 2021). The heightened vulnerability of *T. brucei* to the destruction of the lysosome is demonstrated by the action of the naturally evolved trypanolytic factors TLF-1 and TLF-2 (Vanhamme, 2010). Both factors are high molecular mass human serum components, which cause non-human infectious trypanosomes to lyse. The lytic ingredient in both complexes is apolipoprotein L1 (APOL1). APOL1 is a Bcl-2-like protein that upon acidification forms pores in the lysosomal membrane, which allows the influx of chloride anions and H2O resulting in the uncontrolled osmotic swelling of the lysosome, ultimately destroying the parasite. Disruption of the lysosome as a trypanocidal principle has been demonstrated for a diverse group of reagents, including neuroendocrine peptides (Delgado et al., 2009), L-leucyl-L-leucyl methyl ester (Koh et al., 2015; Alsford, 2015), and different synthetic triterpenoid-peptide conjugates (Leeder et al., 2019). As the second target, we chose the spliced leader (SL)-RNA of *T. brucei*. The 35nt long RNA sequence is the result of a *trans*-splicing reaction and it constitutes the 5’-end of every *T. brucei* mRNA (Günzl, 2010). SL-RNA addition generates monocistronic mRNAs from polycistronic primary transcripts and provides every transcript with a 5’-Cap structure. As such, the reaction is required for the formation of translation-competent mRNAs and, therefore, it is essential. This is further evidenced by the fact that extreme stress conditions result in a complete shutdown of SL-RNA transcription, a phenomenon known as spliced leader silencing (SLS) (Goldshmidt et al., 2010, Menna-Barreto, 2010). Moreover, acridine-derivatized SL-RNA-specific antisense oligodeoxynucleotides have been shown to kill *T. brucei* in cell culture (Cornelissen et al., 1986; Verspieren et al., 1987).

**Figure 1.**
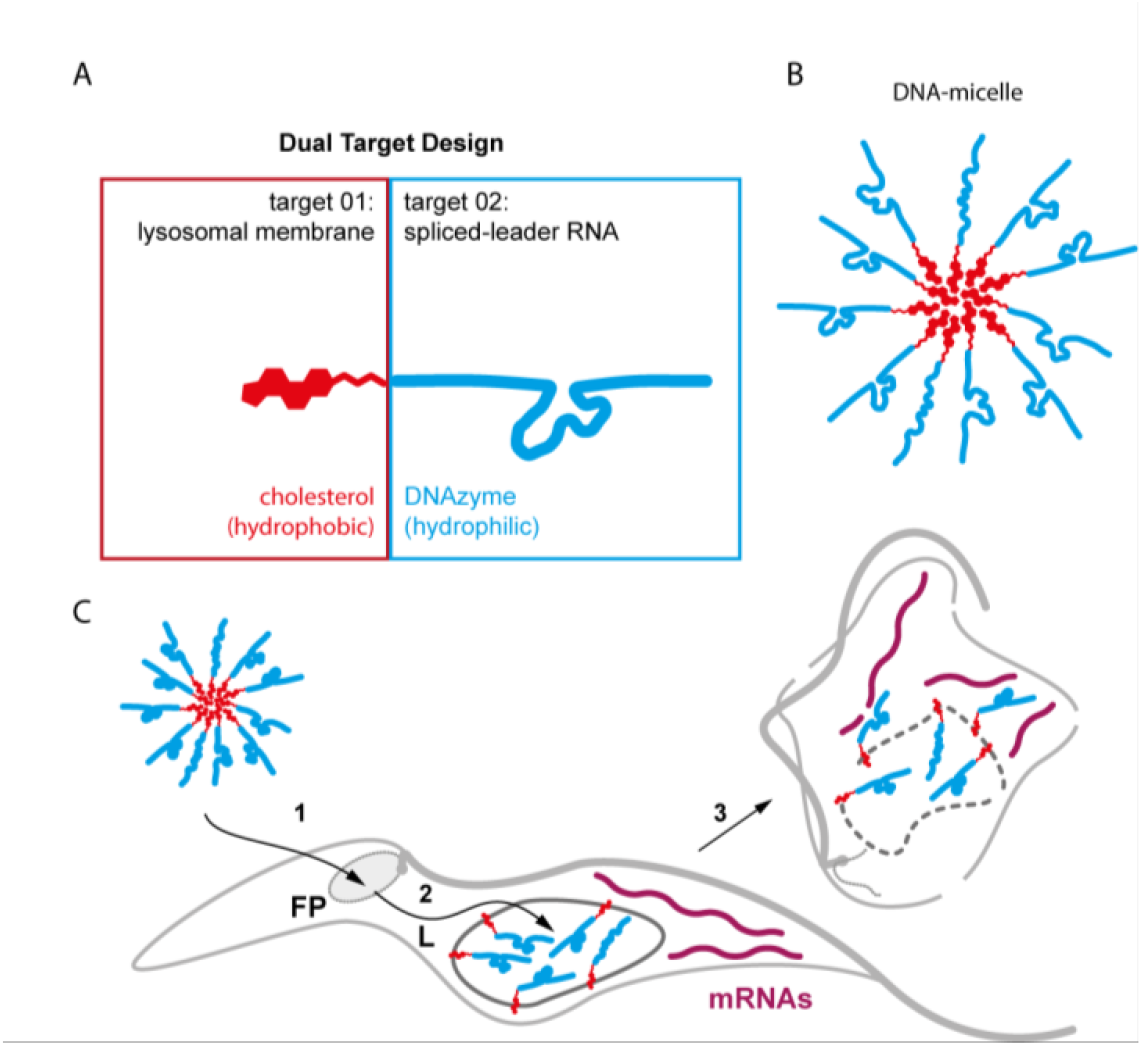
Design of core-shell forming DNA-cholesterol conjugates targeting two parasite-specific pathways (dual-target approach). (A) Bipartite domain structure of a cholesterol-modified oligodeoxyribonucleotide that targets the lysosome (target-1) and the spliced leader (SL)-RNA (target-2) of African trypanosomes. Target-1 is tackled by the cholesterol moiety (sterol scaffold in red) and target-2 is attacked by an RNA-cleaving DNAzyme (cyan). (B) Core-shell nanoparticles self-assemble in aqueous solution resulting in micelle-type high-molecular-mass particles. Red: hydrophobic cholesterol core. Cyan: hydrophilic DNAzyme shell. (C) Putative uptake pathway of the DNAzyme-cholesterol nanoparticles by infective stage trypanosomes. 1: Binding and internalization through the flagellar pocket (FP). 2: Endosomal routing to the lysosome (L). 3: Interaction with the inner lysosomal membrane causes the lysosome to collapse. Escaping DNAzyme-cholesterol molecules catalyze the breakdown of cytosolic mRNAs (purple).

For the synthesis of a lysosomolytic and at the same time SL-RNA-cleaving reagent, we utilize the well-established technology of core-shell DNA-lipid nanostructures (Langecker et al. 2014; Li et al., 2020; Walczak et al., 2021). Derivatizing oligodeoxynucleotides with hydrophobic side chains such as alkyl groups, tocopherol, porphyrins, or cholesterol generates DNA-amphiphiles that self-assemble into macromolecular micelle-type particles. DNA-lipid nanoparticles consist of a membrane-adhesive core and a hydrophilic DNA shell (Figure 1A,B), which are capable of binding and disrupting lipid bilayers. They have been used for pore formation and drug delivery purposes (Huo et al., 2019; Li et al., 2020; Zhao et al., 2021), and specifically cholesterol-derivatized DNA nanoparticles have been applied to functionalize the surface of liposomes, form membrane-spanning nanopores, and induce membrane curvature and tubulations (Singh et al., 2021). The main advantage of DNA-lipid nanoparticles is their programmability. The molecules can be engineered for specific applications by using DNA sequences that execute defined functions (Langecker et al. 2014). Here, we report the synthesis of DNA-lipid nanoparticles consisting of a SL-RNA-specific deoxyribozyme (DNAzyme) covalently attached to a cholesterol group. The construct challenges two essential parasite-specific pathways: the spliced leader sequence at the 5’-end of every *T. brucei* mRNA and the membrane integrity of the parasite lysosome (Figure 1C). We demonstrate nanoparticle formation of the synthesized conjugates and verify that the DNAzyme is enzymatically active within the micellar context. Upon incubation with infective-stage trypanosomes, we show uptake and routing of the particles to the lysosome followed by lysosomal collapse and cell death. As such, we provide evidence for the forward design of a synthetic DNA-lipid nanostructure with therapeutic potential for the treatment of African trypanosomiasis.

## Results and Discussion

### Synthesis of SL-DNAzyme-cholesterol conjugates

SL-RNA-cleaving DNAzyme-cholesterol conjugates were synthesized by automated oligonucleotide synthesis using standard phosphoramidite chemistry. The DNAzyme is 39nt in length and consists of a 13nt 8-17-type DNAzyme core domain (Cruz et al., 2004; Cepeda-Plaza and Peracchi, 2020) flanked by two 12 and 14nt long single-stranded (ss) extensions. The ss-sequences provide base complementary to the *T. brucei* SL-RNA and position the catalytic core of the DNAzyme to cleave the ribose-phosphate backbone between nucleotides G14 and A15 (Figure 2A). Conjugation of the cholesterol moiety was performed co-synthetically either at the 5’- or 3’-end of the DNAzyme using tetraethylenglycol (TEG) or prolinol (Pro) linker chemistries (Figure 2B). Control oligodeoxynucleotides were synthesized without cholesterol and with 5’-stearyl or 5’-oleate residues replacing cholesterol (Figure 2B). Synthesis products were purified by reversed-phase HPLC to purities ≥97% (Supplementary Figure 1) and were confirmed by mass spectrometry. A complete list of all synthesized oligonucleotide-conjugates is provided in Supplementary Table 1.

**Figure 2.**
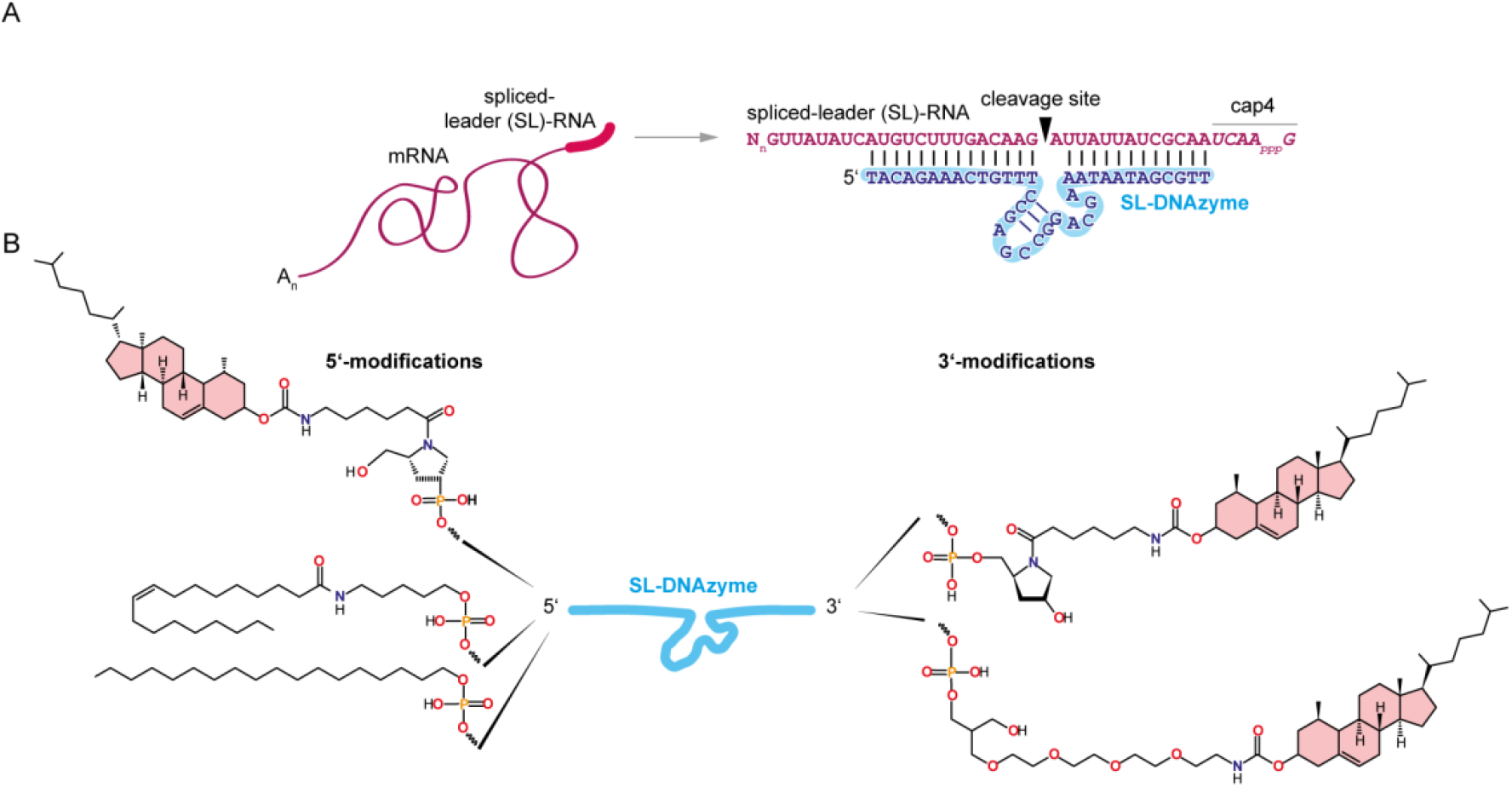
(A) Left: Sketch of a *trans*-spliced and poly-adenylated *T. brucei* mRNA (purple). Right: Nucleotide sequence and base-pairing of a spliced-leader (SL)-RNA/DNAzyme hybrid. Purple: SL-RNA. Cyan: SL-DNAzyme. The SL-RNA is 35nt long and carries a hypermethylated cap4-structure. The sicille phosphodiester bond is marked by an arrowhead. (B) Summary of the different end-modifications of the SL-DNAzyme. 5’-end modifications (top to bottom): prolinol (Pro)-cholesterol, C_6_-oleate, stearyl. 3’-end modifications (top to bottom): Pro-cholesterol, tetraethylene glycol (TEG)-cholesterol. The cholesterol scaffold is colored in red.

### Formation of high molecular mass core-shell nanoparticles

DNA-amphiphiles in aqueous solution self-assemble into high molecular mass core-shell particles (Langecker et al. 2014; Li et al., 2020). The reaction is entropically driven and was experimentally confirmed by isokinetic ultracentrifugation in glycerol gradients. Figure 3A shows a representative result using the SL-DNAzyme 3’-cholesterol conjugate. The molecule assembles into nanoparticles with an apparent S-value of 13S±3S, which calculates to an apparent molecular mass of approximately 450kDa and an apparent aggregation number (N_agg_) of 39±14 monomers/micelle. Figure 3B shows a visualization of the formed micelles using atomic force microscopy (AFM). The sample shows a monodisperse distribution of globular particles ranging from 13 to 23nm in diameter with a mean of 18nm. A further visualization was achieved by gel electrophoresis in native, high-percentage agarose gels (Figure 3C and Supplementary Figure 2). As expected, the different DNAzyme-cholesterol constructs migrate as high molecular mass complexes with apparent sizes between 400 and 530 kDa. This enumerates to a mean apparent N_agg_ of 44 molecules/micelle confirming the N_agg_-measurement by isokinetic ultracentrifugation. Lastly, we measured the critical micelle concentration (CMC) for each of the DNA-cholesterol constructs. For that, we relied on the solvatochromic properties of the benzophenoxazine dye Nile red (Kurniasih et al., 2014). Figure 3D shows the SL-DNAzyme-3’-cholesterol micelle as an example. The molecule has a CMC of 42nM. The CMCs for all DNA-micelle constructs range from 13 to 116nM with a mean of 71nM, which is in agreement with published values, varying between 10-100nM (Liu et al., 2010; Wang et al., 2016; Zou et al., 2018; Yin et al., 2019; Kim et al., 2021). Importantly, the stearate- and oleate-substituted constructs do not form nanoparticles at concentrations ≤3µM (Supplementary Table 1).

**Figure 3.**
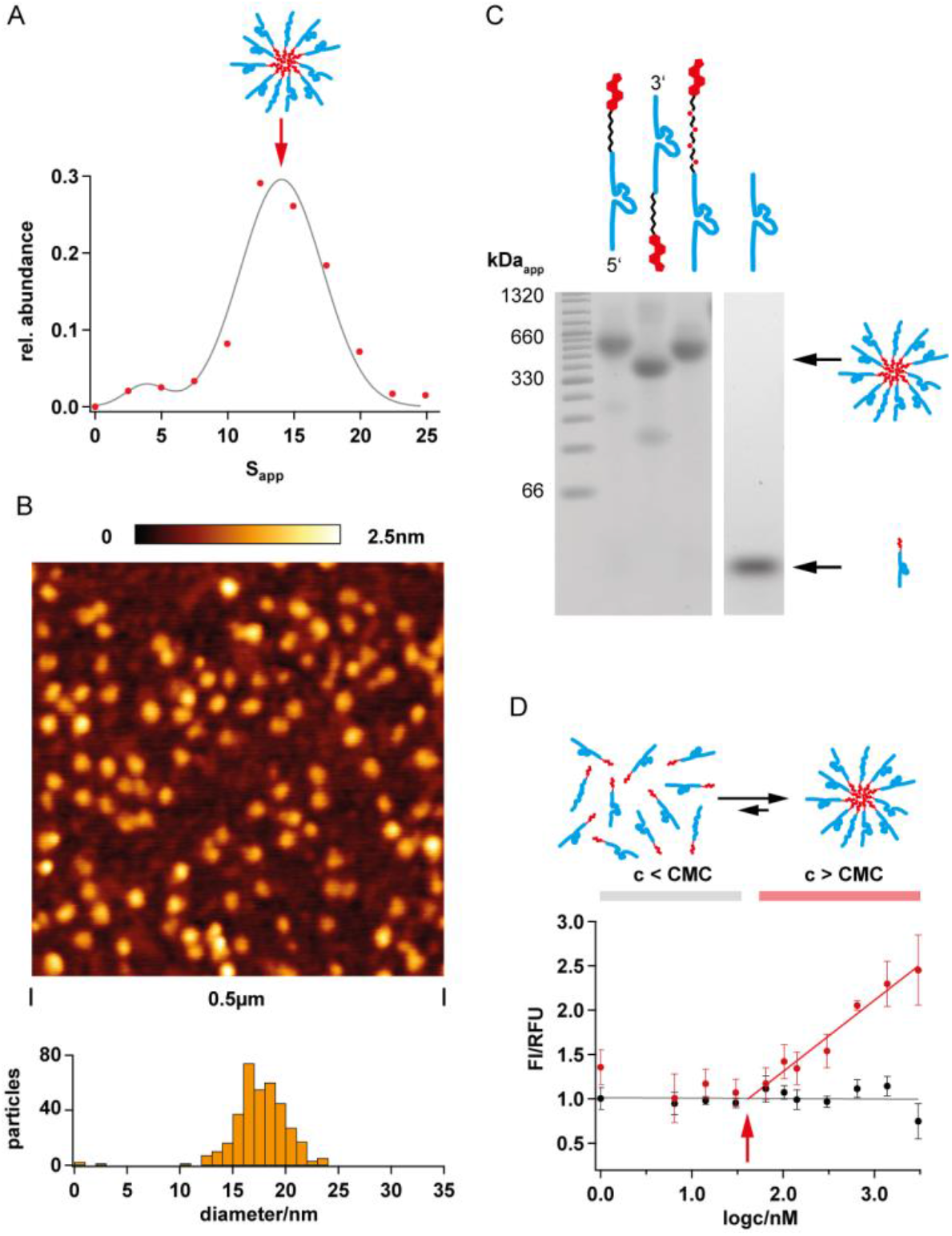
(A) Sedimentation profile of a SL-DNAzyme-cholesterol nanoparticle in a linear 10-35% (v/v) glycerol gradient. The particle sediments with an apparent S-value (S_app_) of 13±3S. (B) Atomic force microscopy (AFM) of SL-DNAzyme-cholesterol nanoparticles on a mica surface. The dried micelles vary in diameter between 13 and 23nm with a mean of 18nm. (C) Electrophoretic separation of different DNAzyme-cholesterol nanoparticles in non-denaturing 3% (w/v) agarose gels (for a summary of the molecule drawings see Supplementary Table 1). The different conjugates migrate with apparent molecular sizes between 400 and 530kDa. (D) CMC measurement of an SL-DNAzyme-cholesterol conjugate using the solvatochromic fluorogenic dye 9-(diethylamino)-5*H*-benzo[*a*]phenoxazin-5-one (Nile red). A list of all measured CMC values is given in Supplementary Table 1. FI= fluorescence intensity. RFU= relative fluorescence unit.

### SL-DNAzyme-cholesterol nanoparticles are catalytically active

The enzymatic activity of the SL-DNAzyme was analyzed *in vitro*. For that, we synthesized a 5’-Cy5-modified SL-DNAzyme and a 5’-FAM-derivatized SL-RNA model substrate. The RNA consists of the first 27nt of the *T. brucei* SL-RNA sequence (Figure 4A) with a single AG-dinucleotide at positions 13/14. Different fluorophores were used for the DNAzyme and the substrate RNA to monitor the two reactants independently by laser-induced fluorescence (LIF). Since the 8-17 DNAzyme is a metalloenzyme, reactions were started by the addition of Mn^2+^. As shown in Figure 4B the SL-oligoribonucleotide is precisely cleaved at the targeted AG-dinucleotide, generating the predicted 13nt long FAM-labeled 5’-cleavage product and as expected for an enzyme, the SL-DNAzyme remains unaltered during the catalytic conversion. Next, we analyzed whether the RNA cleavage reaction can be performed in the context of the DNAzyme-cholesterol nanoparticle (Figure 4C). For that, we used the SL-DNAzyme-3’-cholesterol construct as a representative. Figure 4D shows the cleavage reaction of the SL-oligoribonucleotide at single turnover conditions over a period of 3h. The micelle-embedded DNAzyme cleaves the SL-RNA oligonucleotide with a k_cat_ of 0.01/min and processes up to 75% of the input. Importantly, the nanoparticles are also active under multiple turnover conditions (Figure 4E). At a 10x, 15x, and 30x molar excess of SL-RNA over DNAzyme we measured kcat-values between 0.015 and 0.04/min with up to 95% substrate conversion. Lastly, we determined that the micelle-embedded SL-DNAzyme is active at temperatures between 10°C and 42°C, at pH values between 6.8 to 8.8, and that Mn^2+^ can be replaced by Co^2+^, Ca^2+^, and Mg^2+^ (Supplementary Figure 3).

**Figure 4.**
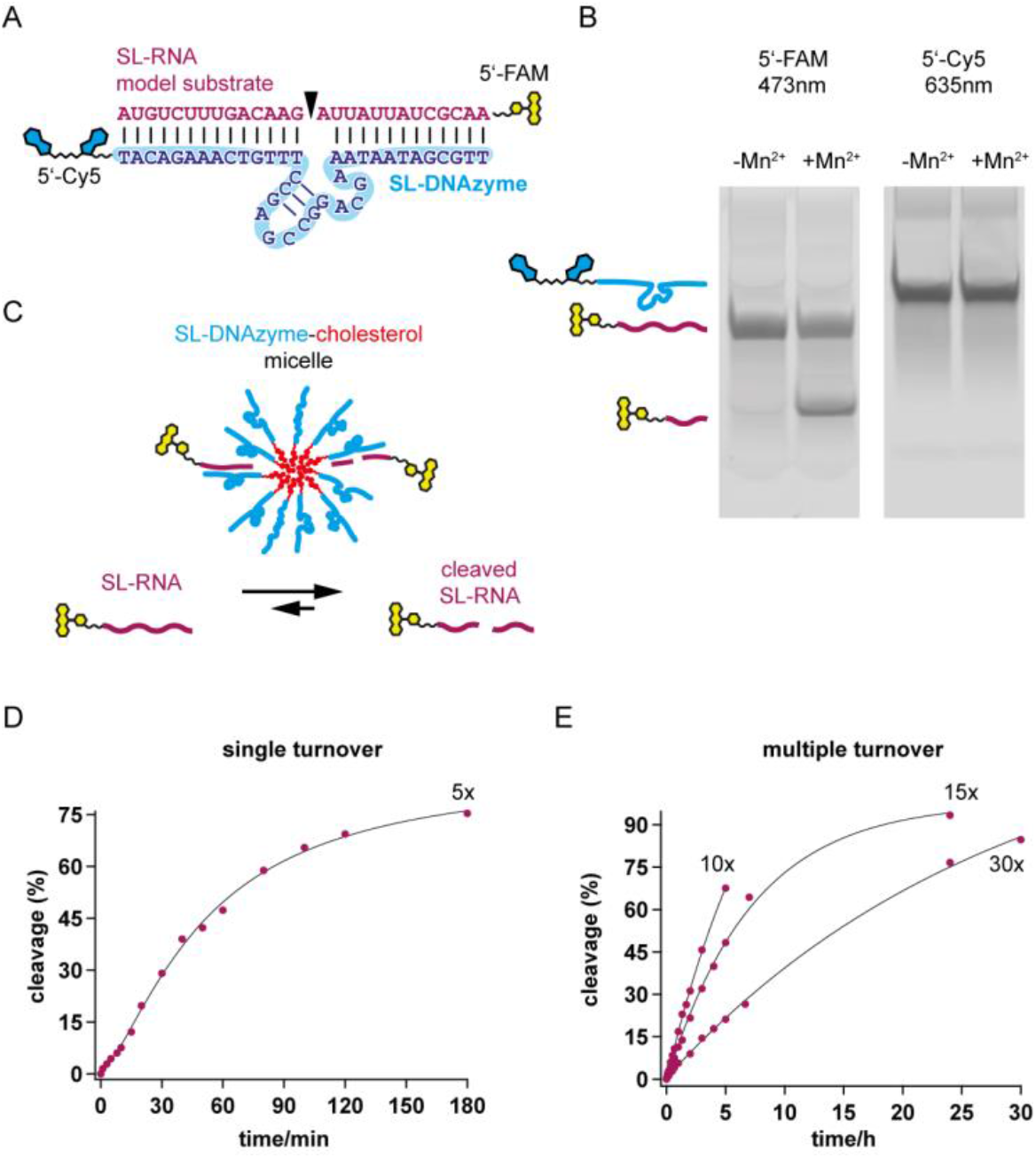
(A) Sequence outline of the SL-DNAzyme/SL-RNA enzyme-substrate complex (DNAzyme=cyan; SL-RNA=purple). The SL-RNA mimicking oligoribonucleotide is 5’-FAM-labeled (fluorescein scaffold in yellow) and the SL-DNAzyme is 5’-Cy5-modified (cyanine scaffold in blue). The catalytic center of the DNAzyme consists of a short hairpin next to four single-stranded nt (ACGA). The RNA cleavage site is an AG-dinucleotide (arrowhead). (B) Representative gel-electrophoretic result of an RNA-cleavage assay in the presence (+) and absence (-) of Mn^2+^. The input SL-RNA oligonucleotide, the 5’-cleavage product, and the SL-DNAzyme are separated in a denaturing polyacrylamide gel followed by LIF-detection at 473nm (FAM) and 635nm (Cy5). The cleavage reaction is Mn^2+^-dependent; as expected, the DNAzyme leaves the reaction unaltered. (C) Graphical outline of the DNAzyme-mediated SL-RNA cleavage reaction in the context of the assembled core-shell nanoparticle. (D) Single turnover kinetic of a SL-RNA cleavage reaction by a SL-DNAzyme-cholesterol nanoparticle at a 5x molar excess of SL-DNAzyme over SL-RNA substrate. Up to 75% of the input SL-RNA is cleaved after 3 hours. (E) The same analysis as in (D) under multiple turnover conditions (10x, 15x, 30x molar excess of SL-RNA over SL-DNAzyme). RNA substrate conversions of up to 90% are achieved.

### SL-DNAzyme cholesterol nanoparticles kill bloodstream-stage trypanosomes

The trypanocidal activity of the different SL-DNAzyme-cholesterol micelles was tested by incubating infective-stage trypanosomes with increasing concentrations (0-320nM) of the nanoparticle constructs. Parasite survival was measured in a live cell assay, based on the cleavage of the membrane-penetrating, fluorogenic calcein-acetoxymethyl (AM) ester by intracellular esterases (Weston and Parish, 1990; Bratosin et al., 2005). As shown in Figures 5A, and B (and Supplementary Figure 4), all DNA-micelle variants act as potent trypanocidals. Half-maximal lethal concentrations (LC_50_) range from 33 to 79nM for the different constructs (Figure 5B) and the trypanocidal activity is neither influenced by the linker chemistry (prolinol or TEG) nor by the 5’- or 3’-positioning of the cholesterol moiety. The killing reaction (at 5xLC_50_) reaches >95% completion after 13 hours, which roughly represents two parasite cell doublings (Figure 5C). Importantly, the unmodified SL-DNAzyme shows no inhibitory activity even in the presence of saturating amounts (67nM) of free cholesterol (Figure 5A) (Saad and Higuchi, 1965). Similarly, the DNA-oleate and DNA-stearate constructs do not have trypanocidal activity (Figure 5B). Also, insect-stage trypanosomes were shown to be unaffected even at concentrations ≥5-fold the LC50. This suggests a specificity for the infective, bloodstream-stage stage of the parasite.

**Figure 5.**
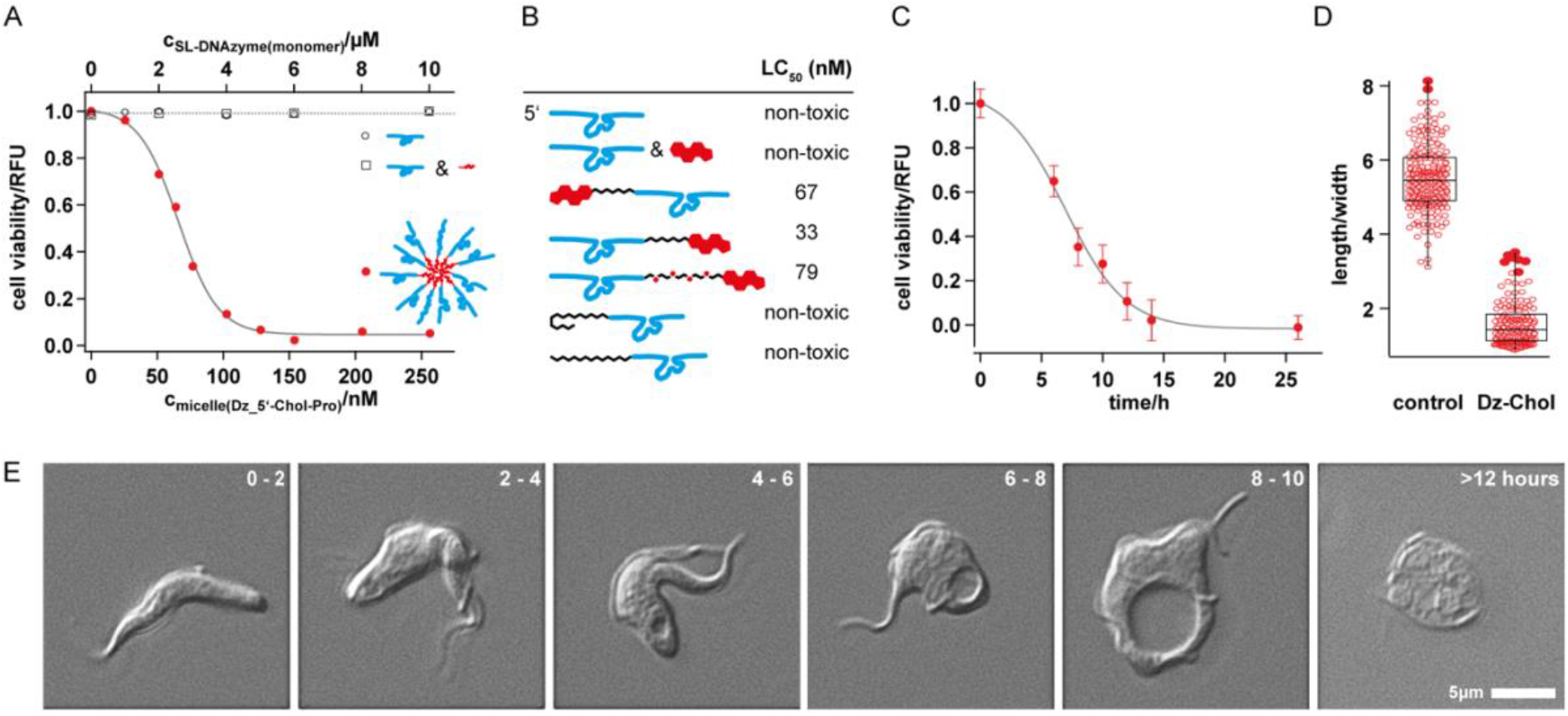
Cell viability analysis. (A) Dose-response curves of infective-stage trypanosomes treated with SL-DNAzyme-cholesterol nanoparticles (filled circles, red), non-conjugated SL-DNAzyme (open circles, black), and non-conjugated SL-DNAzyme in the presence of saturating amounts of free cholesterol (open squares, black). RFU=relative fluorescence unit. (B) Summary of the derived half-maximal lethal concentrations (LC_50_) for the different nanoparticles. Top to bottom: non-conjugated SL-DNAzyme, non-conjugated SL-DNAzyme in the presence of saturating amounts of cholesterol, 5’-Chol-pro-SL-DNAzyme, SL-DNAzyme-3’-pro-Chol, SL-DNAzyme-3’-TEG-Chol, 5’-oleate-SL-DNAzyme, and 5’-stearyl-SL-DNAzyme. For a summary of the molecule drawings see Supplementary Table 1. (C) Cell viability kinetic of infective-stage trypanosomes treated with SL-DNAzyme-cholesterol nanoparticles (8x LC_50_). The half-maximal survival time is 7.0±0.8h. Errors are standard deviations. (D) Box plot analysis of the length/width ratio of infective-stage trypanosomes treated with non-conjugated SL-DNAzyme (c=10µM) and SL-DNAzyme-cholesterol nanoparticles (DzChol c=8x LC_50_). The median length/width ratio changes from 5.4 to 1.4. Incubation time=8h. (E) Representative DIC microscopy images of infective-stage trypanosomes treated with SL-DNAzyme-cholesterol nanoparticles (8x LC_5_) for 0 to >12h as indicated. For a time-lapse movie of the process, see Supplementary Movie 1.

Figure 5E and Supplementary Video 1 show differential interference contrast (DIC) microscopic images of individual DNAzyme-cholesterol nanoparticle-treated *T. brucei* cells. From 2h of incubation, the parasites lose their characteristic slender shape and adopt an enlarged and bloated morphology. This coincides with the appearance of a vacuolar substructure, which increases until it dominates the cell body at 8-10h of incubation. At this stage, the median length/width ratio of the cells has changed from 5.6 to 1.4 (Figure 5D). Further incubation (>12h) results in a complete loss of the trypanosome cell shape, the appearance of intracellular granules, and finally cell lysis. Using FITC-labelled dextran as a fluid phase marker, we identified the vacuolar substructure as the lysosome. FITC-dextran is endocytotically taken up and routed to the lysosome (Ohkuma et al., 1982). As shown in Figure 6A, parasites treated with DNAzyme nanoparticles (for 6-8h) show a perfect overlap of the fluorescent FITC-dextran signal in the lysosome with the large vacuole. This was further confirmed by directly labeling the DNAzyme micelles with 6-carboxyfluorescein (6-FAM) (Figure 6B). As before, at early incubation times (2-4h) the 6-FAM signal perfectly localizes to the lysosome. At 8-10h the signal in the lysosomal lumen diminishes and staining of both, the lysosomal membrane and the cytosol is detected. At 10-12h, the lysosomal signal disappears and spreads out through the entire cell, indicating the collapse of the lysosomal membrane. At >12h, the cells have lost all subcellular structure, and fluorescence is distributed throughout the cytosol, with some punctate signals likely representing stained membrane fragments. Notably, the nucleus and kinetoplast of the nanoparticle-treated parasites are not affected (Figure 6A, B).

**Figure 6.**
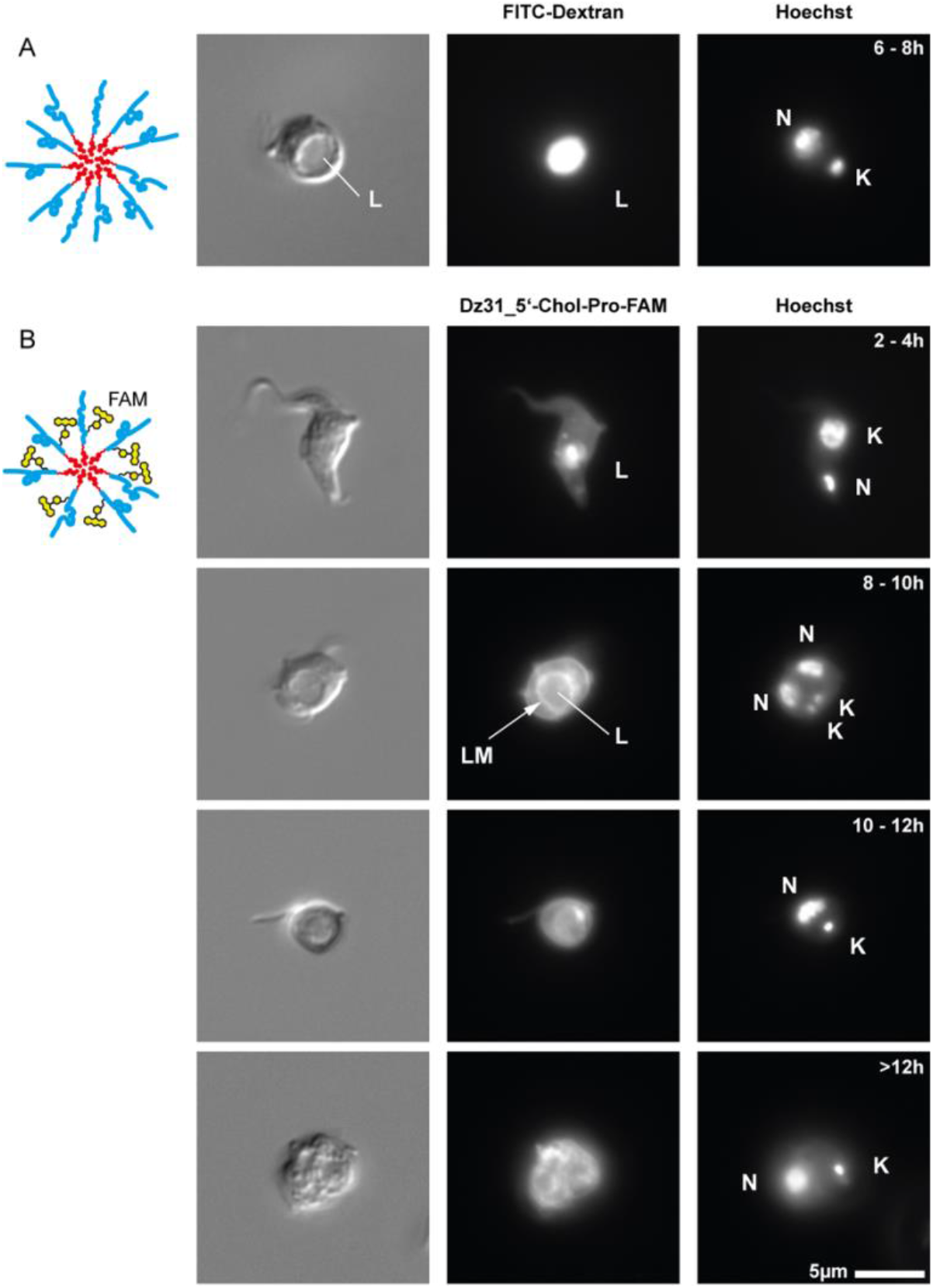
Intracellular localization and fate of the SL-DNAzyme-cholesterol nanoparticles. Left panels: Differential interference contrast (DIC) imaging. Center panels: Fluorescence imaging (FITC and FAM). Right panels: DNA staining. N=nuclear DNA. K=kinetoplast DNA. (A) Visualization of the lysosome (L) using FITC-labeled dextran. (B) Intracellular localization of a FAM-labeled SL-DNAzyme-cholesterol nanoparticle (FAM-DzChol) between 2 and >12h of incubation. LM=lysosomal membrane.

### A functional DNAzyme is not required for the trypanocidal effect

Lastly, we tested whether the trypanolytic activity of the DNAzyme nanoparticles is driven by the cholesterol-mediated collapse of the lysosome, or by the DNAzyme-guided cleavage of the SL-RNA sequence, or both. For that, we synthesized a series of nanoparticle constructs with shortened DNAzyme sequences 31, 20, 10, and 6nt in length. Except for the 31nt long DNA, none of the truncated sequences can fold into a catalytically active DNAzyme. Nevertheless, all constructs show trypanolytic activity (Figure 7A and Supplementary Table 1), proving that the enzymatic activity of the DNAzyme is neither necessary nor sufficient for the trypanolytic effect. LC_50_-values range from 62 to 182nM with an inverse linear relationship between nt-length and LC_50_ (Figure 7B). Further support was gained from the synthesis of a DNA nanoparticle with a randomized 39nt non-DNAzyme sequence. As anticipated the construct displays an LC_50_ of 51nM (Figure 7), almost identical to the 39nt SL-DNAzyme nanoparticles (LC_50_= 33-67nM). Lastly, we established that RNA-based nanoparticles, as well as nuclease-resistant, phosphorothioate-modified DNA sequences, function as trypanocides (Supplementary Table 1). This demonstrates a high sequence and length plasticity of the nucleic acid domain of the nanoparticles (for a summary see Supplementary Figure 5).

**Figure 7.**
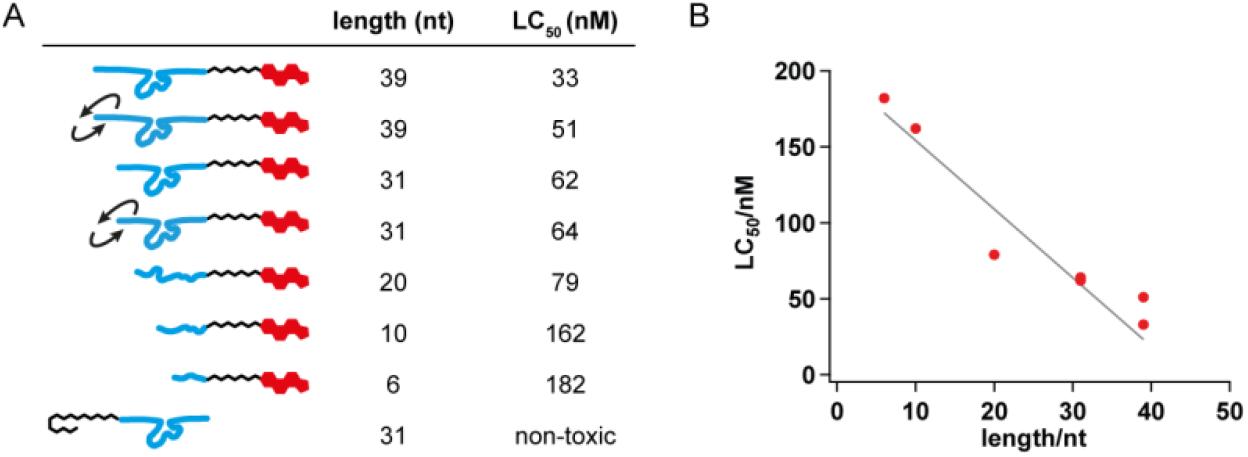
Nucleotide length analysis. (A) LC_50_-values of different DNA-cholesterol constructs ranging from 39nt to 6nt. The different conjugates are depicted as cartoons with the DNA domain in cyan and the cholesterol moiety in red. For a summary of the molecule drawings see Supplementary Table 1. DNA molecules <31nt cannot adopt a functional DNAzyme fold. (B) Nucleotide length-dependency of the measured LC_50_-values. The data are correlated with a Pearson’s *r* = -0.965.

## Conclusions

In summary, we have demonstrated the rational design of nanostructured DNA-cholesterol amphiphiles with trypanocidal activity. The molecules are specific for the infective lifecycle stage of trypanosomes and kill the parasite at nanomolar concentrations (30nM). At this concentration, the amphiphiles form high molecular mass micelles (450kDa, 13S), which are characterized by a hydrophilic solvent-facing DNA shell and a hydrophobic cholesterol core. The nanoparticles are taken up by the parasites, and they are transported to the lysosome, where they execute a membrane-destabilizing effect. As designed, this leads to the collapse of the lysosome, generating an autophagic-like phenotype that ultimately causes cell death. We show that only covalently connected DNA-cholesterol compounds execute the lysosomolytic activity. Neither DNA alone nor cholesterol alone nor a mixture of both molecules is cytotoxic. Furthermore, the trypanocidal effect is strictly dependent on the cholesterol core of the nanoparticles. Cholesterol cannot be substituted by other fatty acid chains, such as oleate or stearate, which do not show inhibitory potency. By contrast, the DNA shell exhibits relaxed characteristics. Phosphorthioate modifications are tolerated, and DNA can be replaced by RNA. Similarly, the length requirement of the DNA shell is relaxed. DNA sequences with a length of 39nt show the highest lysosomolytic potency, however, oligodeoxynucleotides as short as 6nt are still functional. This indicates a sequence-independent role for the DNA shell in the process, possibly contributing to the parasite binding and internalization steps. This is supported by the fact that trypanosomes are deficient in the *de novo* synthesis of purines. The parasites rely on the receptor-mediated uptake of free purines and purine nucleosides (De Konig et al., 2005; Campagnaro et al, 2018), and short DNA and RNA oligonucleotides have also been shown to be internalized (Cornelissen et al., 1886; Verspieren et al., 1987; Homann and Göringer, 1999; Homann and Göringer, 2001; Adler et al., 2008). As such it is feasible that the DNA shell, at least in part, acts as the nanoparticle haptophore. However, the DNA shell also adds to the toxophore properties of the constructs. DNA-cholesterol nanoparticles have been shown to adsorb to lipid membranes by ionic contacts (McManus et al., 2003; Michanek et al., 2009). This is mediated by the negative net charge of the DNA shell, which as a result guides the cholesterol core to the lysosomal membrane (Figure 8). Cholesterol molecules become integrated into the inner leaflet of the lysosomal membrane (Langecker et al., 2014; Morzy et al., 2021; Singh et al., 2021), which generates a highly asymmetric lipid bilayer and as a consequence skews the biomechanical properties of the membrane (Gracia et al., 2010; Subczynski et al., 2017; Chakraborty et al., 2020) (Figure 8). The massive enlargement of the lysosome and the subsequent osmotic swelling of the parasite cell are testament of the compromised membrane integrity (Biswas et al., 2019; Zakany et al., 2020) as is the observed leakage of FITC-dextran from the lysosome into the cytosol after >8h of incubation. Ultimately, as designed, large-scale lysosomal membrane failure, fragmentation of the organelle itself, and fenestration of other membrane compartments kill the parasite. As such, the mode of action (MOA) is qualitatively similar to lysosome-associated defects caused by the natural trypanolytic serum factors TLF1 and TLF2 (Vanhollebeke et al., 2007, Vanwalleghem et al., 2015) and the synthetic trypanolytic factor synTLF (Leeder et al., 2019). Quantitatively, however, DNA-cholesterol nanoparticles exhibit half-maximal lethal concentrations as low as 0.033µM, which is about one order of magnitude lower compared to recombinant APOL1 (0.23µM) (Vanwalleghem, 2015) and about 100 times lower than synTLF (3µM) (Leeder et al., 2019). Lastly, it is important to note that the in-built DNAzyme activity of the nanoparticles, targeting the 5’-end of every *T. brucei* mRNA is dispensable for the trypanocidal activity. DNA-cholesterol nanoparticles with truncated or non-DNAzyme sequences elicit the same cytotoxicity as particles that include the DNAzyme sequence. This establishes that only the membrane fenestration step is necessary and sufficient to cause the trypanocidal effect. However, we cannot exclude that the DNAzyme activity (if present) is masked by the dominant outcome of the lysosomal membrane collapse, and in that case, the SL-DNAzyme activity can provide a backup MOA to counteract the recovery of compromised parasite cells.

**Figure 8.**
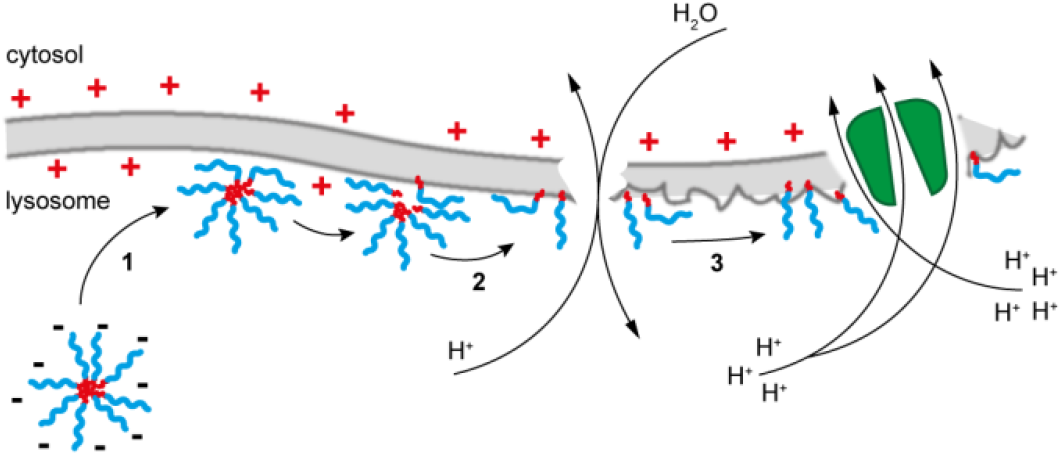
Mechanistic model of the DNA nanoparticle/lipid membrane interaction inside the *T. brucei* lysosome. Step 1: Electrostatic binding of the polyanionic DNA shell to the inner lysosomal membrane. Step 2: Destabilization of the DNA shell and membrane integration of the hydrophobic cholesterol molecules. Step 3: The accumulation of cholesterol molecules in the luminal leaflet of the lysosome generates an asymmetric/dysfunctional membrane architecture. This causes the leakage of protons into the cytosol and the influx of water into the lysosome. Lysosomal swelling and disruption of the membrane ultimately kill the parasite.

## Experimental Section

### Growth of trypanosome cells

The bloodstream life cycle stage of *Trypanosome brucei brucei* strain Lister 427 (MITat serodeme, variant clone MITat 1.2) was grown in HMI-9 medium (Hirumi and Hirumi, 1989) supplemented with 10% (v/v) heat-inactivated fetal calf serum (FCS), 0.2mM 2-mercaptoethanol and 100U/mL penicillin/streptomycin (Gibco™ Thermo Fisher Scientific). Parasites were grown at 37° C in 95% air, and 5% CO_2_ at a relative humidity (rH) ≥95%. Insect-stage (procyclic) parasites were grown at 27°C in SDM-79 medium as described in Brun and Schönenberger, 1979. Parasite cell densities were determined by automated cell counting.

### Synthesis of cholesterol-modified DNA-oligonucleotides

Cholesterol-modified oligodeoxynucleotides were synthesized by automated solid-state synthesis on controlled pore glass (CPG)-beads (200nmol synthesis scale) using 5’-Dimethoxytrityl-2’-deoxy,3’-[(2-cyanoethyl)-(N,N-diisopropyl)]-phosphoramidites. Cholesteryl moieties were co-synthetically introduced either at the 5’- or 3-ends of the different oligonucleotides using triethylene glycol (TEG) or prolinol (Pro) linker chemistries. Control oligonucleotides were synthesized with 5’-stearyl or 5’-oleate modifications, with backbone phosphorothioate (PTO) modifications, and with a 6-Carboxyfluorescein (6-FAM) fluorescence tag. In addition, an “all-RNA” variant was synthesized using 2’-O-*tert*-butyldimethylsilyl (TBDMS)- protected phosphoramidites. The crude synthesis products were purified by reverse phase high-performance liquid chromatography (RP-HPLC), analyzed by mass spectrometry (MALDI-TOF), and further scrutinized in 8M urea-containing, 18% (w/v) polyacrylamide gels (Supplementary Figure S1). Concentrations were derived from UV-absorbency measurements at 260nm (A_260_) using the molar extinction coefficients (ε in L/mol x cm) listed below. The following oligonucleotide sequences were synthesized:

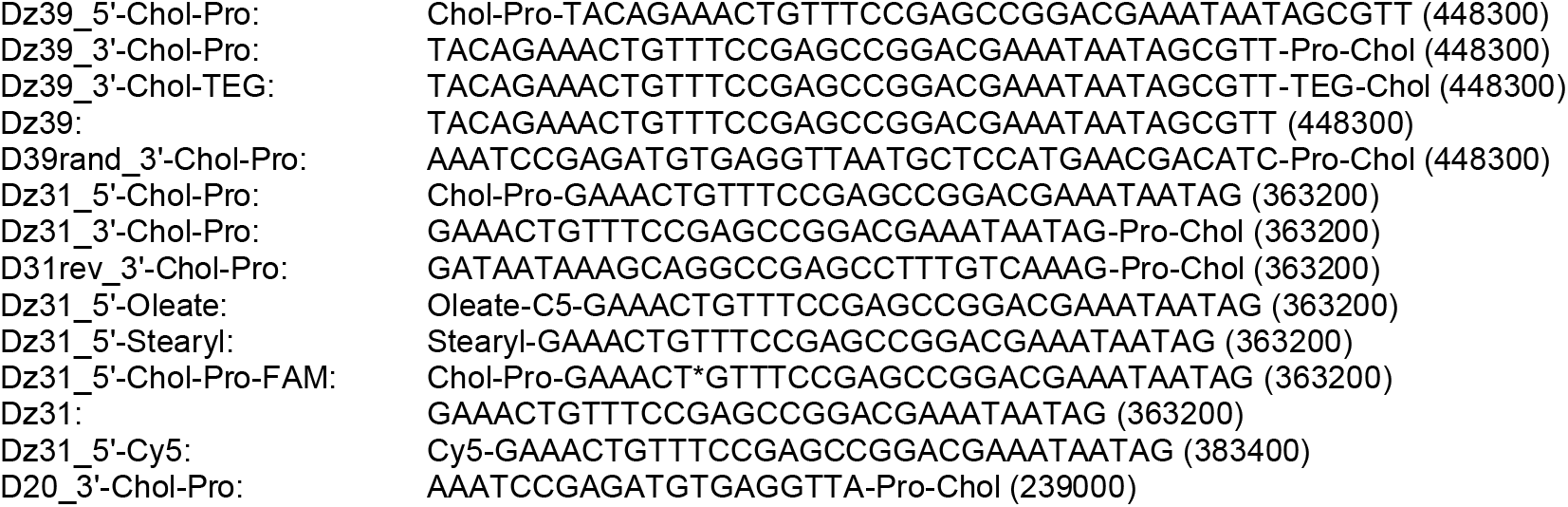
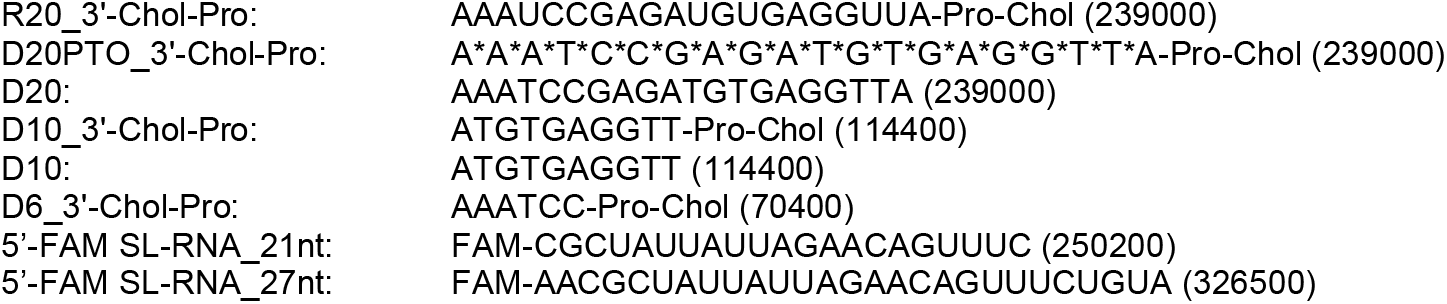

### Formation of DNA-cholesterol nanoparticles and CMC-determination

Cholesterol-modified oligonucleotides were solubilized in 10mM Tris/HCl pH 7.5, 1mM EDTA (TE)-buffer at a concentration of 0.1mM. When aqueous buffer is added, micelles are formed by spontaneous self-association. Critical micelle concentrations (CMC) were determined as in Kurniasih et al., 2015. For that the cholesterol-modified oligodeoxynucleotides were taken up in 25mM 9-(diethylamino)-5*H*-benzo[*a*]phenoxazin-5-one (Nile red, C_20_H_18_N_2_O_2_) in anhydrous MeCN. Samples were incubated at 20°C overnight followed by the stepwise addition of phosphate-buffered saline (PBS) pH7.4 to a final DNA-cholesterol concentration of 3µM. Dilutions for fluorescence measurements were made using freshly prepared 0.03mM Nile red in PBS and measurements were performed in transparent, flat-bottom 96-well plates in a final volume of 0.1mL. Fluorescence spectra (λ_ex_ 535nm) were recorded in 2nm steps in a TECAN Infinite 200PRO plate-reader. Relative fluorescent units (RFU) were integrated between 630 and 680nm and the fluorescence intensities were normalized and plotted as a function of the concentration of DNA-cholesterol. Data points were subjected to a linear fit to calculate the CMC.

### Sedimentation analysis of DNA-cholesterol nanoparticles

DNA-cholesterol micelles at ≥10xCMC were formed in 20mM HEPES/KOH pH7.5, 10mM MgCl_2_, 0.1mM Na_2_EDTA containing either 150mM NaCl or 30mM KCl. Samples were loaded onto 2 mL linear 10-35% (v/v) glycerol gradients and centrifuged at 20°C for 155min at 45.000rpm in a Beckmann Coulter TLS55-rotor. Gradient fractions (0.2mL) were EtOH-precipitated and the DNA-cholesterol material in each fraction was analyzed in denaturing (8M urea), 18% (w/v) polyacrylamide gels. The gels were stained with Toluidine Blue O and densitometrically analyzed. Apparent sedimentation coefficients (S_app_) and the corresponding apparent molecular masses (MW_app_) of the nanoparticles were calculated from a non-linear fit of the sedimentation behavior of the following size standards: *E. coli* tRNA (4S, 2.8×10^4^ Da), *E. coli* 5S rRNA (5S, 4.2×10^4^ Da), *E. coli* 16S rRNA (16S, 5.4×10^5^ Da), *E. coli* 23S rRNA (23S, 1×10^6^ Da), *T. brucei* CR4 pre-mRNA (9S, 1.6×10^5^ Da) and *T. brucei* Cyb pre-mRNA (14S, 4.3×10^5^ Da). Nanoparticle aggregation numbers (N_agg_) were calculated as N_agg_=MW_micelle_/MW_monomer_.

### Visualization of DNA-cholesterol nanoparticles

DNA-cholesterol micelles were electrophoretically separated in 3% (w/v) agarose gels in TBE pH8.3 buffer (89mM Tris-OH, 89mM B(OH)_3,_ 2mM EDTA) followed by staining with the fluorescent cyanine dye SYBR-gold (λ_ex_ 473nm, λ_em_ 510). Stained gels were densitometrically analyzed using a FLA-5000 Imager (FUJIFILM) and quantified with the help of FIJI (Schindelin et al., 2019) and Igor Pro 6.37 (Wave Metrics). Further visualization was achieved by atomic force microscopy (AFM). DNA-cholesterol micelles (10µM) in PBS containing 5mM MgCl_2_ were deposited on freshly cleaved muscovite mica (Plano GmbH) and incubated for 5-10min at RT. The samples were rinsed with dH_2_O and dried in a gentle N_2_ stream. Microscopy was performed using a NanoWizard3 Atomic Force Microscope (JPK) equipped with a 160AC-NA (µmasch-Europe) cantilever (ν=300kHz, D=26N/m). Radial profiles of the micelles were extracted using Gwyddion 2.58 (Nečas and Klapetek, 2012) and Gaussian fits of the profiles were used to approximate the full width at half maximum (FWHM).

### DNAzyme characterization

DNAzyme-driven cleavage reactions were performed in 50mM Tris/HCl pH7.6, 150mM NaCl at a 5’-FAM-labelled SL-RNA substrate concentration of 5µM. The RNA was mixed with a 5-fold molar excess of 5’-Cy5-labelled SL-DNAzyme and samples were denatured for 1min at 75°C. Formation of the SL-RNA/DNAzyme complex was achieved by cooling to 20°C at a rate of 0.08°C/s. Following hybridization, samples were thermally equilibrated at 37°C and cleavage reactions were started by adding 10mM MnCl_2_. After 30min, samples were put on ice and supplemented with 50mM Na_2_EDTA pH8. Reaction products were analyzed in 8M urea-containing 18% (w/v) polyacrylamide gels and analyzed by fluorescence densitometry using the following wavelength: λ_ex_ 473nm, λ_em_ LP510nm, and λ_ex_ 635nm, λ_em_ LP665). Fluorescence signals were quantified using MultiGauge v3.0 (FUJIFILM). Single-turnover enzyme reactions were performed as above using 2.5µM 5’-FAM-labelled SL-RNA and a 5-fold molar excess of 3’-cholesterol modified SL-DNAzyme. Samples were incubated at 37°C in the presence of 10mM MgCl_2_. The divalent cation-dependency of the cleavage reaction was measured at single-turnover conditions in the presence of 10mM of either BaCl_2_, CaCl_2_, CoCl_2_, CuCl_2_, MgCl_2_, MnCl_2_, NiCl_2_, SrCl_2_, or ZnCl_2_. The pH-optimum was analyzed in 25mM Tris/HCl between pH6.8 and pH8.8. Multiple-turnover kinetics were measured similarly using 15µM 5’-FAM SL-RNA and 0.5µM, 1.0µM, or 1.5µM 3’-cholesterol modified SL-DNAzyme. The turnover number (k_cat_) was enumerated from the linear part of the kinetic where ≤30% of the RNA substrate is processed.

### Trypanosome growth inhibition

Growth inhibition experiments were performed in 96-well plates using mid- to late-log phase trypanosomes at a cell density of 0.45×10^6^/mL. 2×10^4^ parasites in 0.05mL of HMI-9 medium were supplemented with 1-10µM DNA-cholesterol nanoparticles and were grown for 16-46h at 37°C. After incubation 5µM of the cell-permeant compound, Calcein AM (C_30_H_26_N_2_O_13_, Life Technologies) was added and the cells were further incubated for 30min at 37°C. Cells were pelleted (2000x g, 10min, 4°C), washed with ice-cold PBSG (PBS plus 5% (w/v) glucose), and resuspended in 0.2mL PBSG. Samples were transferred into transparent, flat-bottom 96-well plates and analyzed using a FLA-5000 imager (FUJIFILM) (λ_ex_ 473nm, λ_em_ 510nm). Fluorescent signals were quantified using FIJI (Schindelin et al., 2019). Cells treated with 10% (v/v) EtOH were used for background subtraction and replicates of the background-subtracted samples were averaged and normalized to the maximal signal. LC_50_-values were calculated as the concentration at 50% fluorescence intensity, using the built-in sigmoidal fit function of Igor Pro 6.37 (Wave Metrics).

### Microscopy and live-cell imaging

Mid-log bloodstream stage trypanosomes were harvested by centrifugation (3000x g, 2min, RT) and resuspended in HMI-9 medium to a cell density of 0.8×10^6^/mL. The parasites were supplemented with 10µM DNA-cholesterol nanoparticles and 5mg/mL fluorescein isothiocyanate (FITC)-dextran with a mean molecular mass of 10kDa (Sigma-Aldrich) as a fluid phase marker. Cells were grown at 37°C for 8-16h followed by pelleting at 2000x g for 5min at 4°C. Cells were washed with ice-cold PBSG and fixed with 1% (w/v) paraformaldehyde and 0.08% (w/v) glutaraldehyde in PBS. Fixed cells were supplemented with 1µg/mL Hoechst 33258 (C_25_H_37_Cl_3_N_6_O_6_, Sigma-Aldrich) and imaged directly or in the presence of 0.5 vol. DABCO antifade (20mM 1,4-diazabicyclo[2.2.2]octane in 90% (v/v) glycerol). Imaging was performed with a Zeiss Axioskop 2 microscope equipped with a Zeiss Axiocam MRm camera and a Plan Neofluar 100x (NA 1.30) oil-objective. Zeiss filter sets for Hoechst 33258 (λ_ex_ BP365/12, λ_em_ LP397) and FITC/FAM (λ_ex_ BP470/20, λ_em_ BP505-530) were used. Cell shape measurements were made after 8h of incubation with 10µM DNA-cholesterol nanoparticles. Cells were fixed and imaged as above, except using a Plan Neofluar 40x air-objective (NA 0.75). Cell diameters were measured using FIJI (Schindelin et al., 2019). For life-cell imaging, trypanosomes were harvested by centrifugation at 3000x g for 5min at 4°C. Cells were resuspended in HMI-9 medium to 5×10^6^ cells/mL and supplemented with 20µM DNA-cholesterol nanoparticles (∼500x CMC and 15x LC_50_). The samples were incubated at 37°C and aliquots were taken at 4h, 6h, 8h, 10h, and 14h and imaged directly using a Plan Neofluar 100x (NA 1.30) oil-objective in DIC-mode using an Axiocam MRm camera at 16 frames/sec. Images were cropped and normalized by dividing the individual frames by the median pixel intensity using FIJI (Schindelin et al., 2019).

## Acknowledgments

*The authors thank Michael Brecht and Jacqueline Nentwich for their experimental help at an early stage of the project. Andreas Völker is thanked for maintaining trypanosome cultures and Elisabeth Kruse for discussions. The work was supported by the Illing Foundation for Molecular Chemistry to HUG*.

