## Supplementary_Information for "Core-shell DNA-cholesterol nanoparticles exert lysosomolytic activity in African trypanosomes"

### Table of Contents

**1 SI Tables and Figures**

1.1 Table S1
1.2 Figure S1
1.3 Figure S2
1.4 Figure S3
1.5 Figure S4
1.6 Figure S5

### 1 SI Tables and Figures

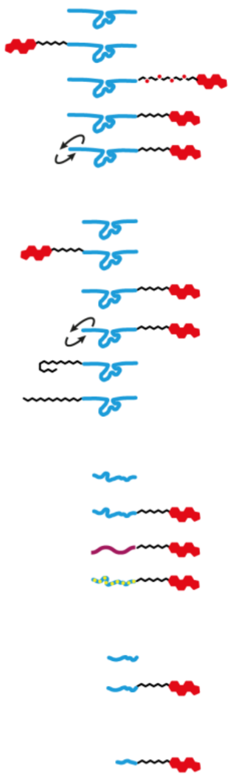

| cartoon depiction | designation | length (nt) | nucleic acid | lipid modification | attachement site | linker chemistry | antiSL-DNAzyme sequence | aggregation number | cmc | $LC_{50}$ |
|---|---|---|---|---|---|---|---|---|---|---|
| | Dz39 | 39 | DNA | none | none | none | + | none | none | non-toxic |
| | Dz39_5'-Chol-Pro | 39 | DNA | cholesterol | 5' | pro | + | n.a. | 13 | 67 |
| | Dz39_3'-Chol-TEG | 39 | DNA | cholesterol | 3' | TEG | + | 28 | 116 | 79 |
| | Dz39_3'-Chol-Pro | 39 | DNA | cholesterol | 3' | pro | + | 33 | 42 | 33 |
| | D39rand_3'-Chol-Pro | 39 | DNA | cholesterol | 3' | pro | - | n.a. | 112 | 51 |
| | Dz31 | 31 | DNA | none | none | none | + | none | none | non-toxic |
| | Dz31_5'-Chol-Pro | 31 | DNA | cholesterol | 5' | pro | + | 30 | 46 | 79 |
| | Dz31_3'-Chol-Pro | 31 | DNA | cholesterol | 3' | pro | + | 69 | 148 | 62 |
| | D31rev_3'-Chol-Pro | 31 | DNA | cholesterol | 3' | pro | - | n.a. | 42 | 64 |
| | Dz31_5'-Oleate | 31 | DNA | oleate | 5' | pro | + | n.a. | none | non-toxic |
| | Dz31_5'-Stearyl | 31 | DNA | stearyl | 5' | pro | + | 32 | none | non-toxic |
| | D20 | 20 | DNA | none | none | none | - | none | none | non-toxic |
| | D20_3'-Chol-Pro | 20 | DNA | cholesterol | 3' | pro | - | 42 | 112 | 79 |
| | R20_3'-Chol-Pro | 20 | RNA | cholesterol | 3' | pro | - | n.a. | 87 | 178 |
| | D20PTO_3'-Chol-Pro | 20 | PTO | cholesterol | 3' | pro | - | n.a. | 330 | 136 |
| | D10 | 10 | DNA | none | none | none | - | none | none | non-toxic |
| | D10_3'-Chol-Pro | 10 | DNA | cholesterol | 3' | pro | - | 39 | 151 | 162 |
| | D6_3'Chol-Pro | 6 | DNA | cholesterol | 3' | pro | - | 30 | 94 | 182 |

**Table S1.** Cartoon representation and data summary of all synthesized oligonucleotides and DNA/RNA-lipid conjugates. DNA and RNA sequences are depicted as lines in cyan (DNA), purple (RNA) and green/yellow (phosphorothioate (PTO)-modified DNA). The presence of the SL-DNAzyme is indicated by an outline of the 2D structure. Cholesterol modifications are in red. The individual sequences of the different constructs are listed in the experimental section of the manuscript. Table S1 includes the micellar aggregation number, the critical micelle concentration (CMC), and the half-maximal lethal concentration ($LC_{50}$) for infective stage trypanosomes. TEG =triethylene glycol. pro=prolinol.

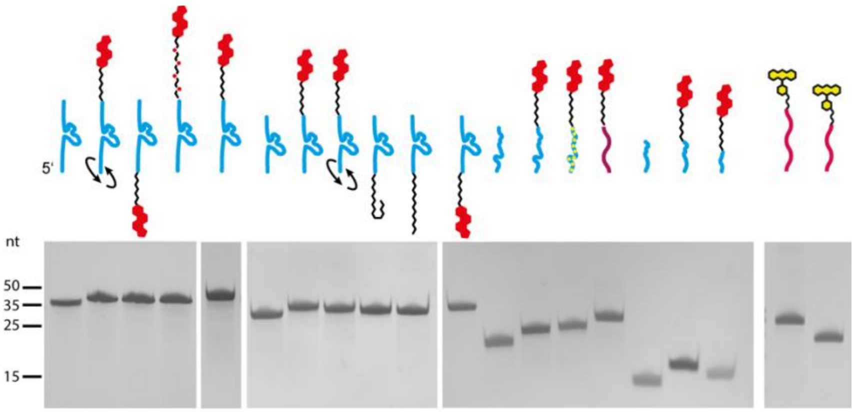


**Figure S1.** Quality control of chemical synthesis. Gel electrophoretic separation of all synthesized DNA-lipid conjugates, RNA-lipid conjugates, and SL-RNA-mimicking oligoribonucleotides in urea-containing (8M) 18% (w/v) polyacrylamide gels. Purities are ≥97%. For a summary of the molecule drawings see Supplementary Table S1.

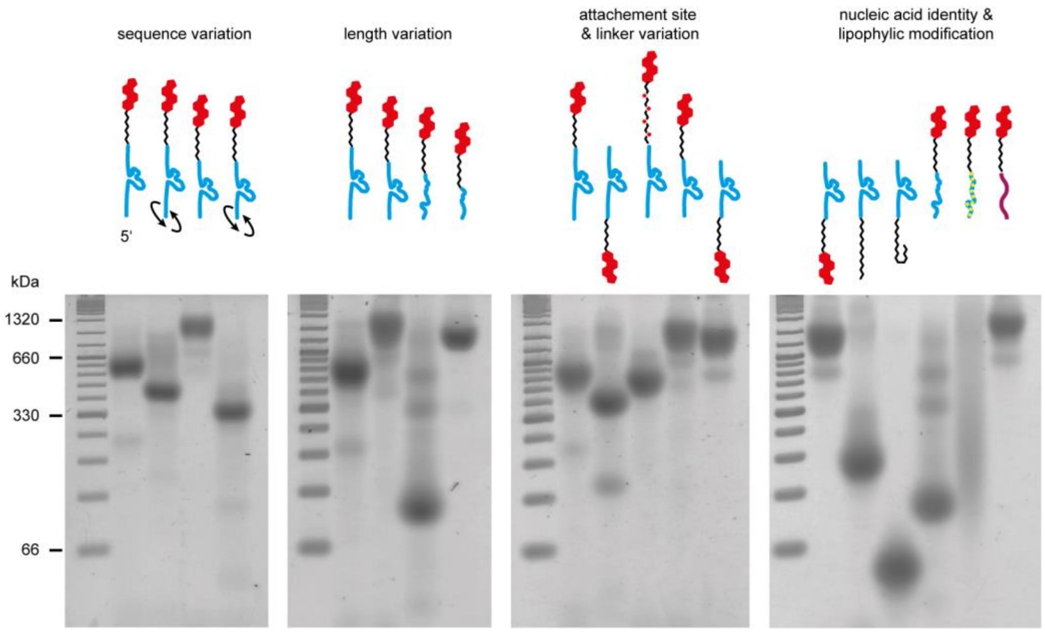


**Figure S2.** High molecular mass nanoparticle formation of different DNA- and RNA-lipid conjugates analyzed in non-denaturing 3% (w/v) agarose gels. The different constructs are grouped (left to right) according to their sequence and length variation, the 5'- or 3'- attachment of the cholesterol moiety, the variation in linker chemistry (prolinol *versus* TEG), the nucleic acid identity (DNA, RNA, PTO-modified DNA) and the lipohilic side-group (cholesterol, oleate, stearate). For a summary of the molecule drawings see Supplementary Table S1.

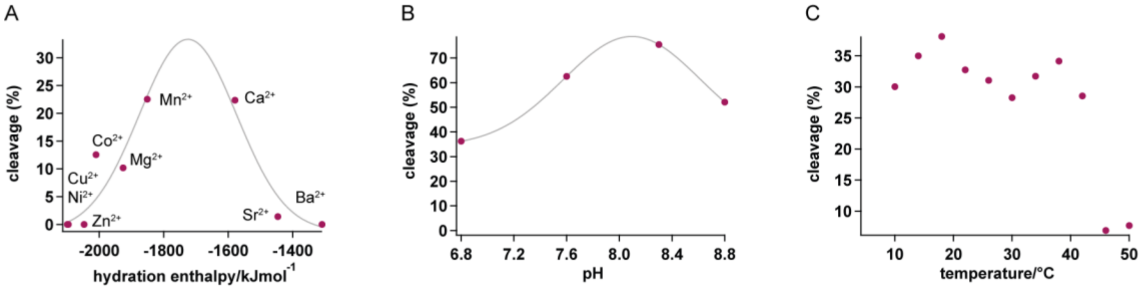


**Figure S3.** Characteristic features of the SL-RNA-specific DNAzyme. (A) Divalent cation dependency. (B) pH-optimum. (C) Temperature optimum.

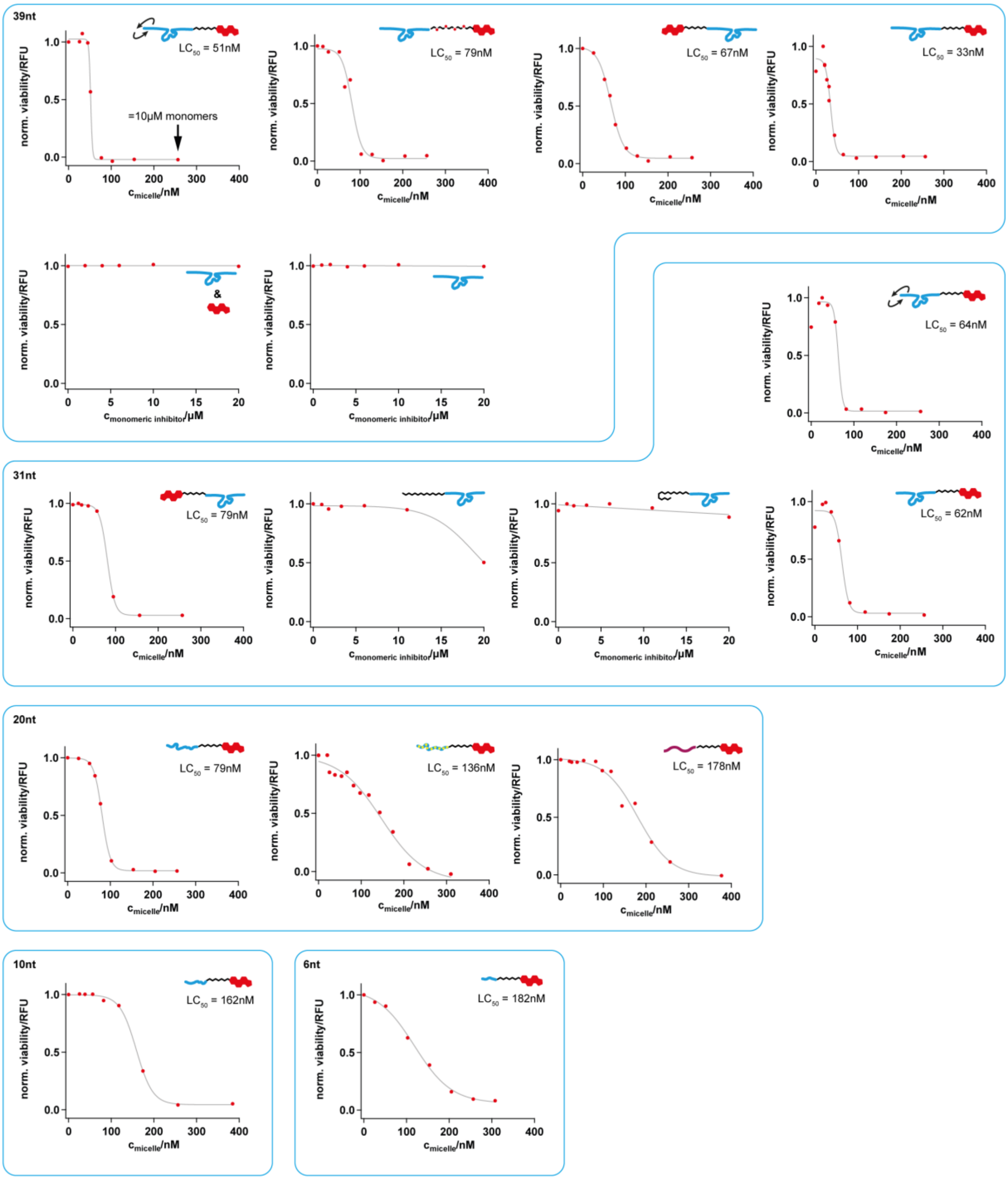


**Figure S4.** Dose-response curves and $LC_{50}$-values of infective-stage trypanosomes treated with different DNA-lipid nanoparticles. The various plots are grouped according to the nucleotide length of the DNA/RNA domain of the nanoparticle constructs. Molecule drawings are listed in Supplementary Table S1. RFU=relative fluorescence unit.

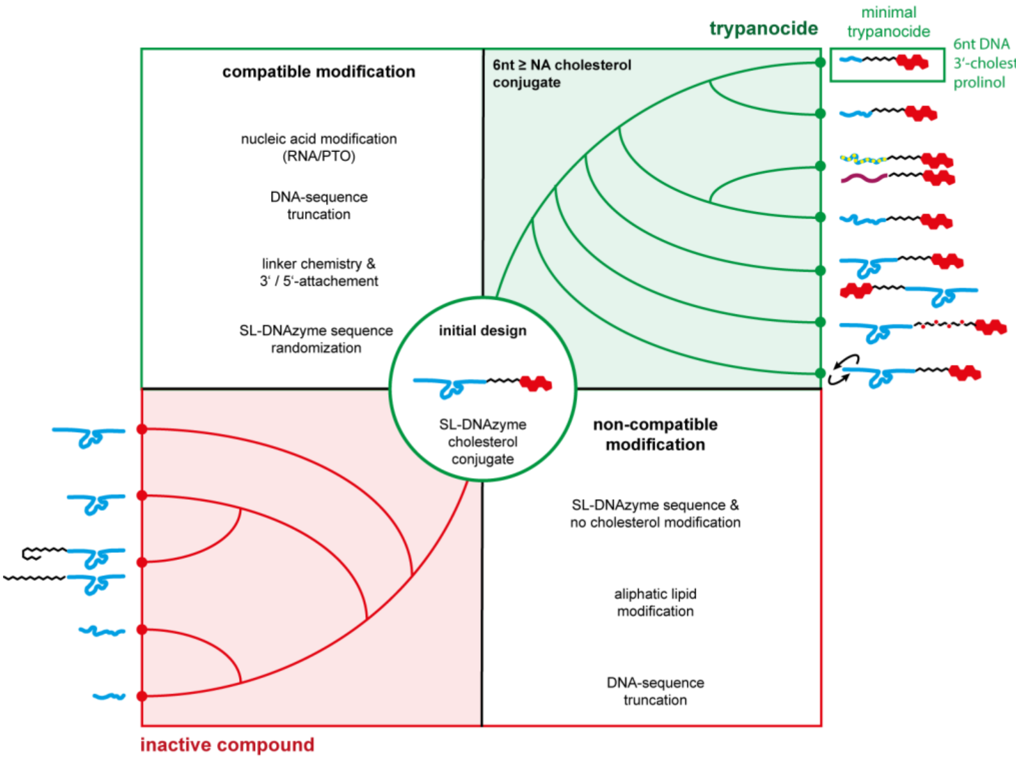


**Figure S5.** Summary of compatible and non-compatible modifications and characteristics for the synthesis of DNA-cholesterol nanoparticles with trypanocidal activity.
